# Acute head shaking precedes chronic corpus callosum deficits during repeated cocaine exposure in common marmosets

**DOI:** 10.64898/2026.09.11.749290

**Authors:** Sun Mi Gu, Chul Kyu Lee, Taeyun Yoo, Jae Jun Lee, Heejong Eom, Jeong Pyo Son, Tae Hwan Kim, Chun-Woong Park, Seong Shoon Yoon, Dohyun Lee, Jaesuk Yun

**Affiliations:** College of Pharmacy, Chungbuk National University, 194-31 Osongsaengmyeong 1-ro, Osong-eup, Heungdeok-gu, Cheongju-si, Chungcheongbuk-do 28160, Republic of Korea; Non-clinical Evaluation Center, Osong Medical Innovation Foundation, 123 Osongsaengmyeong-ro, Osong-eup, Heungdeok-gu, Cheongju-si, Chungcheongbuk-do 28160, Republic of Korea; College of Pharmacy, Catholic University of Daegu, 13-13 Hayang-ro, Hayang-eup, Gyeongsan-si, Gyeongsangbuk-do 38430, Republic of Korea; College of Korean Medicine, Daegu Haany University, 136 Sincheondong-ro, Suseong-gu, Daegu 42158, Republic of Korea; Accelerator Radioisotope Research Section, Advanced Radiation Technology Institute (ARTI), Korea Atomic EnergyResearch Institute (KAERI), 29 Geumgu-gil, Jeongeup-si, Jeonbuk-do 56212, Republic of Korea

**Keywords:** Cocaine, Marmoset, Artificial intelligence, Head shaking, Brain connectivity, Diffusion tensor imaging, DPYSL2, CRYAB

## Abstract

**Background:** Stimulant exposure can induce acute stereotyped behaviors and chronic alterations in brain connectivity and white matter integrity. However, the temporal sequence and molecular changes linking these cocaine-induced phenotypes remain unclear.

**Methods:** We combined LabGym-based behavioral analysis, longitudinal brain imaging, and cross-species molecular analysis in 11 common marmosets (*Callithrix jacchus*; seven male, four female): four underwent behavioral and imaging studies, six provided molecular data, and one provided immunohistochemical data. Male *Cryab* knockout and wild-type mice underwent functional studies.

**Results:** Acute cocaine (5 mg/kg, intraperitoneally) induced rapid, repetitive lateral head movements, defined as head shaking. After 1 and 10 months of repeated exposure, resting-state functional connectivity was altered in sensory, parietal, prefrontal, motor, and hippocampal regions. Changes were detected at 1 month, whereas diffusion tensor imaging showed reduced fractional anisotropy and axial diffusivity in the corpus callosum splenium at 10 months, with no significant genu changes. Cross-species transcriptomic and proteomic comparison identified DPYSL2 and DNM3 as shared axon-associated molecules. Western blotting showed reduced DPYSL2 in brain tissue from cocaine-treated marmosets. CRYAB localized to O4-positive callosal oligodendrocytes and increased following chronic cocaine exposure. *Cryab* knockout mice had reduced corpus callosum thickness, increased forced-swim immobility, and reduced open-field distance traveled, supporting a role for CRYAB in corpus callosum integrity and depression-related behavior.

**Conclusions:** These findings show that acute cocaine-induced stereotyped head shaking precedes functional connectivity changes and later corpus callosum deficits. Reduced DPYSL2 may be associated with axonal dysfunction, whereas increased CRYAB may represent a response to cocaine-induced white matter stress.

## Introduction

Cocaine causes acute neurobehavioral effects and persistent changes in brain function. In nonhuman primates (NHPs), psychostimulants can induce neuropsychiatric symptoms and stereotyped repetitive movements (1). Prolonged cocaine exposure disrupts circuits involved in reward processing, impulse control, and cognition (2,3) and induces structural brain changes and cognitive deficits in rhesus macaques (4). However, the temporal and molecular links between acute cocaine-induced behaviors and chronic neural abnormalities remain unclear.

Neuroimaging studies identify functional and structural connectivity abnormalities in cocaine use disorder. Altered thalamocortical connectivity suggests impaired cortical-subcortical communication (5). Diffusion tensor imaging (DTI) studies also show reduced fractional anisotropy (FA) in chronic cocaine users, including interhemispheric white matter pathways, and associate reduced white matter integrity with poor treatment outcomes (6). The corpus callosum, the largest interhemispheric white matter tract, integrates distributed brain networks; its disruption may therefore contribute to persistent connectivity and behavioral abnormalities.

White matter integrity depends on axonal, oligodendrocytic, cytoskeletal, and cellular stress-response processes that chronic cocaine exposure may disrupt. Integrating human postmortem transcriptomic data with proteomic data from experimentally exposed NHPs can identify molecular candidates associated with axonal and callosal abnormalities; functional studies are then needed to determine their roles.

The common marmoset (*Callithrix jacchus*) is well suited to studying cocaine-related brain and behavioral changes because of its human-like neuroanatomy and compatibility with neuroimaging. Our previous marmoset studies examined cocaine self-administration with behavioral and imaging changes (7) and 2C-B reinforcement with dopamine transporter binding (8). Here, LabGym quantified head shaking after acute cocaine administration; resting-state functional magnetic resonance imaging and DTI assessed functional connectivity and white matter integrity after 1 and 10 months of repeated exposure. We integrated our published human nucleus accumbens transcriptomic data (9) with marmoset brain proteomics to identify axon-associated candidates, validated selected proteins in the marmoset brain, and tested a corpus callosum-associated factor in a genetic mouse model. We aimed to define an acute stereotyped behavior preceding chronic brain-network and callosal abnormalities and identify associated molecular factors.

## Methods and Materials

### Animals

A total of 11 common marmosets (*Callithrix jacchus*; seven males and four females), aged three years at study onset, were obtained from the Osong Medical Innovation Foundation (Chungbuk, Republic of Korea). Four underwent acute behavioral assessment and longitudinal neuroimaging before cocaine exposure and after 1 and 10 months of repeated administration. An independent cohort of six was divided into control (*n* = 3) and cocaine-treated (*n* = 3) groups for brain proteomics and Western blot validation; one additional marmoset underwent immunohistochemistry. Marmosets were housed individually in standard cages (45 × 60 × 60 cm) at 27 ± 2°C and 40 ± 10% relative humidity, with 10–15 air changes per hour and a 12-h light/dark cycle (lights on, 07:00–19:00; illumination ≥500 lux). They received a formulated diet (50 g/day; No. 0630, Altromin, Lage, Germany) and sterilized water *ad libitum*. Marmoset experiments were conducted at the foundation’s Laboratory Animal Center with Institutional Animal Care and Use Committee approval (KBIO-IACUC-2020-148). Male *Cryab* knockout (KO) mice on a C57BL/6N background were produced by Macrogen Co., Ltd. (Seoul, Republic of Korea) using CRISPR/Cas9 gene editing, as described previously (9). *Cryab* KO and wild-type (WT) mice underwent behavioral and histological assessment of the corpus callosum. Mouse experiments followed the Guide for the Care and Use of Laboratory Animals and were approved by the Animal Care Committee of Chungbuk National University (Cheongju, Republic of Korea; CBNUA-2027-22-02).

### Cocaine administration and longitudinal imaging schedule

Cocaine hydrochloride (Toprak Mahsulleri Ofisi, Türkiye) was administered intraperitoneally at 5 mg/kg, six days per week, for up to 10 months. The dose was based on a study of repeated intraperitoneal cocaine in marmosets (10) and our previous marmoset cocaine study (11).

Baseline imaging occurred within 10 days before the first cocaine dose; follow-up imaging occurred after 1 and 10 months of repeated exposure. Cocaine was suspended during each follow-up, and scans were completed within 10 days after the last scheduled dose. After the 1-month session, administration resumed on the same schedule until the 10-month assessment.

### AI-based quantification of cocaine-induced head shaking

Before testing, marmosets were moved from catching cages to the experimental room. Sessions occurred between 09:30 and 15:00 in a transparent acrylic chamber approximately 530 mm wide, 445 mm deep, and 410 mm high. A front-mounted digital camera recorded video at 29.97 frames per second and either 1280 × 720 or 1920 × 1080 pixels, depending on the camera, in MP4 format. Within the first 2 weeks of cocaine administration, the four marmosets underwent a 30-min baseline recording followed by 30 min after cocaine (5 mg/kg, intraperitoneally). LabGym classified head shaking, immobility, and low activity in 15-frame windows. Head shaking comprised repeated lateral head movement more than once per window; immobility, no discernible limb or trunk movement; and low activity, detectable low-intensity movement such as slow head or subtle limb movement or brief limited locomotion. The LabGym Categorizer was trained on 4,494 paired animation-pattern examples: 1,624 head-shaking, 1,844 low-activity, and 1,026 immobility examples, each spanning 15 consecutive frames. The pattern-image standard deviation threshold was 50; the Animation Analyzer and Pattern Recognizer each used complexity level 4. Performance was evaluated in a categorizer dataset of 450 pairs (162 head-shaking, 185 low-activity, and 103 immobility) and an independent test categorizer dataset of 224 pairs (80 head-shaking, 93 low-activity, and 51 immobility). Precision, recall, F1-score, and overall accuracy were calculated; both evaluations achieved 0.91 accuracy. Supplementary 4 provides detailed metrics and representative original and AI-annotated videos. A separate video dataset then quantified the three behaviors before and after cocaine in the four marmosets and generated frame-wise raster plots.

### Magnetic resonance imaging (MRI)

During image acquisition, the animals were anesthetized with 1.5% isoflurane delivered through a face cone in a 3:7 N O/O mixture at a flow rate of 1 L/min. Respiratory rate and cardiac pulse were monitored using a small-animal monitoring system (SA Instruments, Inc., Stony Brook, NY, USA). Body temperature was maintained at 36.5 ± 1°C using a water-circulating heating bed and monitored using a rectal temperature probe. All MRI measurements were performed using a Bruker BioSpec 47/40 USR system (Ettlingen, Germany) equipped with a birdcage volume coil with an inner diameter of 72 mm for signal transmission and reception. For cerebral blood volume-weighted resting-state functional magnetic resonance imaging (rs-fMRI), ferumoxytol (Feraheme, AMAG Pharmaceuticals, Lexington, MA, USA) was administered intravenously at 15 mg/kg before image acquisition. CBV-weighted rs-fMRI images were acquired using a gradient-echo echo-planar imaging sequence with the following parameters: repetition time (TR) = 2,000 ms, echo time (TE) = 12 ms, bandwidth = 150 kHz, matrix size = 96 × 60, field of view (FOV) = 40 × 25 mm², slice thickness = 1.2 mm, number of slices = 30, number of repetitions = 300, and total acquisition time = 10 min. For DTI, images were acquired using a pulsed-gradient spin-echo echo-planar imaging sequence with the following parameters: TR = 4,000 ms, TE = 42 ms, bandwidth = 325 kHz, matrix size = 192 × 120, FOV = 40 × 25 mm², slice thickness = 1.0 mm, number of slices = 35, number of averages = 14, 30 diffusion-encoding gradient directions, and five *b* = 0 images.

### ROI-based functional connectivity analysis

The EPI data were preprocessed to improve the detection of resting-state hemodynamic signal fluctuations. Preprocessing included slice-timing correction, temporal despiking, linear detrending, motion correction by realigning each volume to the first volume, band-pass filtering between 0.01 and 0.2 Hz, regression of nuisance variables including motion parameters, and spatial smoothing using a Gaussian kernel with a full width at half maximum of 0.6 mm. Data processing and analysis were performed using Analysis of Functional NeuroImages and MATLAB (MathWorks, Natick, MA, USA). Individual functional images were linearly registered to a customized marmoset EPI template using the FMRIB Linear Image Registration Tool with 12 degrees of freedom, including translation, rotation, scaling, and shearing. Fifteen regions of interest (ROIs), comprising 10 cortical and five additional regions, were defined using the customized EPI template: dorsolateral prefrontal cortex (DLPFC), ventrolateral prefrontal cortex (VLPFC), medial prefrontal cortex (MPFC), orbitofrontal cortex (OFC), motor and premotor cortex (MOT), somatosensory cortex (SS), auditory cortex (AU), posterior parietal cortex (PPC), posterior cingulate and retrosplenial cortex, visual cortex, hippocampal formation (HipF), thalamus (Thal), caudate nucleus (Cd), accumbens nucleus (Acb), and cerebellum (CeB). The locations of the ROIs are shown in Supplementary 1. Resting-state functional connectivity was quantified by calculating correlation coefficients between the signal time courses of each pair of predefined ROIs. Changes in rsFC between baseline and 1 month and between baseline and 10 months after cocaine administration were analyzed using paired Student’s *t*-tests.

### DTI data analysis

DTI preprocessing included correction for eddy-current distortions and head motion. Data processing and diffusion tensor fitting were performed using TrackVis and the FMRIB Software Library. FA and axial diffusivity (AD) were calculated from the eigenvalues of the diffusion tensor. The genu of the corpus callosum (GCC) and splenium of the corpus callosum (SCC) were defined as ROIs. FA and AD values were extracted from each ROI at baseline and at 1 and 10 months after cocaine administration. Because the same four marmosets were examined at all three time points, longitudinal changes in each DTI parameter were analyzed using one-way repeated-measures analysis of variance with Greenhouse–Geisser correction. When a significant effect of time was detected, paired comparisons between time points were performed using paired Student’s *t*-tests with Holm adjustment.

### Proteomics analysis

Proteomic analysis was performed using pooled cerebral cortical tissue containing the underlying white matter, including the corpus callosum, from control (*n* = 3) and 10-month cocaine-treated (*n* = 3) marmosets. Tissue samples were pooled by group, and 50 μg of protein from each pool was analyzed. Samples were denatured by adding 100 μL of 8 M urea (final urea concentration, ≥6 M). Disulfide bonds were reduced with 1 μL of 1 M dithiothreitol during a 40-min incubation at 56 °C in a ThermoMixer C (Eppendorf, Hauppauge, NY, USA). Proteins were then alkylated with 2.5 μL of 1 M iodoacetamide for 40 min at room temperature in the dark. Samples subsequently received 900 μL of 25 mM ammonium bicarbonate and were digested with a trypsin/Lys-C protease mixture (Promega) at an enzyme-to-protein ratio of 1:50. Digestion was performed at 37°C for 16 h using a ThermoMixer C. The resulting peptides were desalted on C18 solid-phase extraction columns, dried, and resuspended in 0.1% formic acid at 0.5 μg/μL for liquid chromatography–tandem mass spectrometry (LC– MS/MS). Peptides were analyzed using an EASY-nLC 1000 system coupled to an EASY-Spray source and a Q Exactive Hybrid Quadrupole-Orbitrap mass spectrometer (Thermo Fisher Scientific). A 2-μL aliquot of each sample was loaded onto a 2-cm Acclaim PepMap 100 C18 trapping column (75 μm inner diameter, 3 μm particle size, 100 Å pore size, nanoViper; Thermo Fisher Scientific). Peptides were separated on a 50-cm EASY-Spray PepMap RSLC C18 analytical column (75 μm inner diameter, 2 μm particle size, 100 Å pore size; Thermo Fisher Scientific) maintained at 50°C. Protein identification and database searches used the Sequest algorithm in Proteome Discoverer version 1.4 (Thermo Fisher Scientific) against the *Callithrix jacchus* proteome database. Detailed LC–MS/MS acquisition parameters are provided in Supplementary 2. For cross-species comparison, we used our previously published RNA-sequencing data from the nucleus accumbens of human drug users (9). Differentially expressed genes in the human dataset were compared with differentially expressed proteins in the marmoset dataset after filtering for molecules associated with the Gene Ontology cellular component term “axon.”

### Western blotting

Cerebral cortical tissue containing the underlying white matter was obtained from the same control (*n* = 3) and 10-month cocaine-treated (*n* = 3) marmosets used for proteomic analysis. Individual tissue samples were used for Western blot validation of selected proteins. Proteins were extracted as previously described (12, 13) using lysis buffer containing 20 mM Tris-HCl, 250 mM NaCl, 2 mM EDTA, and 1% Triton X-100. The primary antibodies used were anti-CRYAB (ab76467, Abcam; 1:1,000), anti-DPYSL2 (CSB-PA614975YA01HU, CUSABIO; 1:1,000), and anti-GAPDH (2118S, Cell Signaling Technology; 1:1,000). HRP-conjugated anti-mouse IgG (RABHRP2-10UL, Sigma-Aldrich; 1:5,000) and HRP-conjugated anti-rabbit IgG (RABHRP1-10UL, Sigma-Aldrich; 1:5,000) were used as secondary antibodies. Protein bands were visualized using an enhanced chemiluminescence reagent and a Fusion Solo S imaging system (Vilber Lourmat, France). Band intensities were quantified using ImageJ software (Wayne Rasband). The intensity of each target protein band was normalized to that of GAPDH, and the normalized values were expressed relative to the mean value of the control group.

### Measurement of plasma cocaine concentration

Approximately 4 h after the final cocaine dose, 1 mL of blood was collected from each of the three cocaine-treated proteomics marmosets into heparinized tubes on ice, centrifuged at 2,000 × *g* for 15 min at 4 °C, and stored as plasma at −80 °C. A SCIEX API 4000 mass spectrometer coupled to an Agilent 1100 HPLC system quantified cocaine by modified published LC–MS/MS methods (14,15). Supplementary 3 provides chromatographic and MS/MS conditions.

### Fluorescent immunohistochemistry

Fluorescent immunohistochemistry was used to localize CRYAB in oligodendrocytes within the marmoset corpus callosum. Brain tissue was fixed with 4% paraformaldehyde and cryoprotected in sucrose solution. Coronal sections (10-μm thick) were washed with phosphate-buffered saline, permeabilized with Triton X-100, and blocked with 3% bovine serum albumin for 1 h at room temperature.

The sections were incubated overnight at 4°C with primary antibodies against CRYAB (ab76467, Abcam; 1:300) and O4 (O7139, Sigma-Aldrich; 1:300). After washing, the sections were incubated for 1 h at room temperature in the dark with goat anti-mouse IgG conjugated to Alexa Fluor 488 (ab150113, Abcam; 1:500), goat anti-rabbit IgG conjugated to Alexa Fluor 488 (ab150077, Abcam; 1:500), goat anti-mouse IgG conjugated to Alexa Fluor 568 (ab175473, Abcam; 1:500), and goat anti-rabbit IgG conjugated to Alexa Fluor 568 (ab175471, Abcam; 1:500). Nuclei were counterstained with DAPI (300 nM; Sigma-Aldrich). Fluorescence images were acquired using an Axio Imager.A2 fluorescence microscope (Carl Zeiss, Oberkochen, Germany). The localization of CRYAB in O4-positive oligodendrocytes was examined in the corpus callosum.

### Behavioral profiling and Klüver–Barrera staining in *Cryab* KO mice

Spontaneous locomotor activity was assessed in 8-week-old male WT (*n* = 7) and *Cryab* KO (*n* = 9) mice. Each mouse was placed individually in the center of a square activity box (30 × 30 × 25 cm) and allowed to move freely for 60 min. Behavior was video-recorded and analyzed with the SMART-LD video-tracking system (Panlab, Barcelona, Spain), and total distance traveled was quantified. The forced swim test was conducted using an independent cohort of 8-week-old male WT and *Cryab* KO mice (*n* = 6 per group). Each mouse was placed individually for 5 min in a cylindrical container (20 cm in diameter and 30 cm high) containing water. Behavior was recorded and analyzed with the SMART-LC video-tracking system (Panlab, Barcelona, Spain), and total immobility duration and percentage of time spent immobile were quantified. For histological assessment of corpus callosum integrity, 10-μm-thick coronal brain sections were prepared from 16-week-old WT and *Cryab* KO mice. Sections were selected at three anteroposterior levels: anterior CC (approximately Bregma +1.1 mm), middle CC (approximately Bregma +0.8 mm), and posterior CC (approximately Bregma +0.5 mm). Sections were rehydrated in 100% and 95% ethanol for 5 min each, then washed with phosphate-buffered saline and distilled water for 2 min each. After incubation in Luxol Fast Blue solution (IW-3005A, IHC World, Woodstock, MD, USA) at 56°C for 16 h, excess stain was removed with 95% ethanol and distilled water. Sections were differentiated in 0.05% lithium carbonate for 30 s and then in 70% ethanol until the gray matter appeared transparent and the white matter was clearly distinguishable. They were subsequently washed with distilled water for 1 min and counterstained with 0.1% cresyl violet for 5 min. Sections were then rinsed with distilled water for 1 min, dehydrated in 95% and 100% ethanol for 1 min each, and cleared in xylene. Finally, they were mounted with Permount (Fisher Scientific, Hampton, NH, USA) and examined at 100× magnification using an Axio Imager A2 light microscope (Carl Zeiss, Oberkochen, Germany).

### Statistical analysis

Data are presented as the mean ± standard error of the mean. Paired Student’s *t*-tests were used to compare acute behavioral measures before and after cocaine administration in the same marmosets. Changes in resting-state functional connectivity between baseline and 1 month and between baseline and 10 months after cocaine administration were also analyzed using paired Student’s *t*-tests. Longitudinal changes in DTI parameters across baseline, 1 month, and 10 months were analyzed using one-way repeated-measures analysis of variance with Greenhouse–Geisser correction. When a significant effect of time was detected, paired comparisons between time points were performed using paired Student’s *t*-tests with Holm adjustment. Unpaired Student’s *t*-tests were used for Western blot comparisons between control and cocaine-treated marmosets and for behavioral comparisons between WT and *Cryab* KO mice. Statistical analyses were performed using SigmaPlot version 14 (Systat Software, San Jose, CA, USA) and SPSS version 18 (SPSS Inc., Chicago, IL, USA). A *P* value < 0.05 was considered statistically significant.

## Results

### AI-based behavioral analysis identifies cocaine-induced stereotyped head shaking

The LabGym Categorizer distinguished head shaking, low activity, and immobility with 0.91 overall accuracy in both the performance and independent Test Categorizer datasets; Supplementary 4 provides detailed performance and representative original and AI-annotated videos. Raster plots showed infrequent pre-cocaine head shaking that increased after acute cocaine (Fig. 1). Counts increased in all four marmosets from 246.5 ± 78.5 before cocaine to 833.3 ± 241.9 afterward (paired Student’s *t*-test, *P* = 0.039). Low-activity counts also increased (*P* < 0.05), whereas immobility decreased nonsignificantly. Thus, AI-based video analysis quantified acute cocaine-induced stereotyped head shaking.

**Fig. 1.**
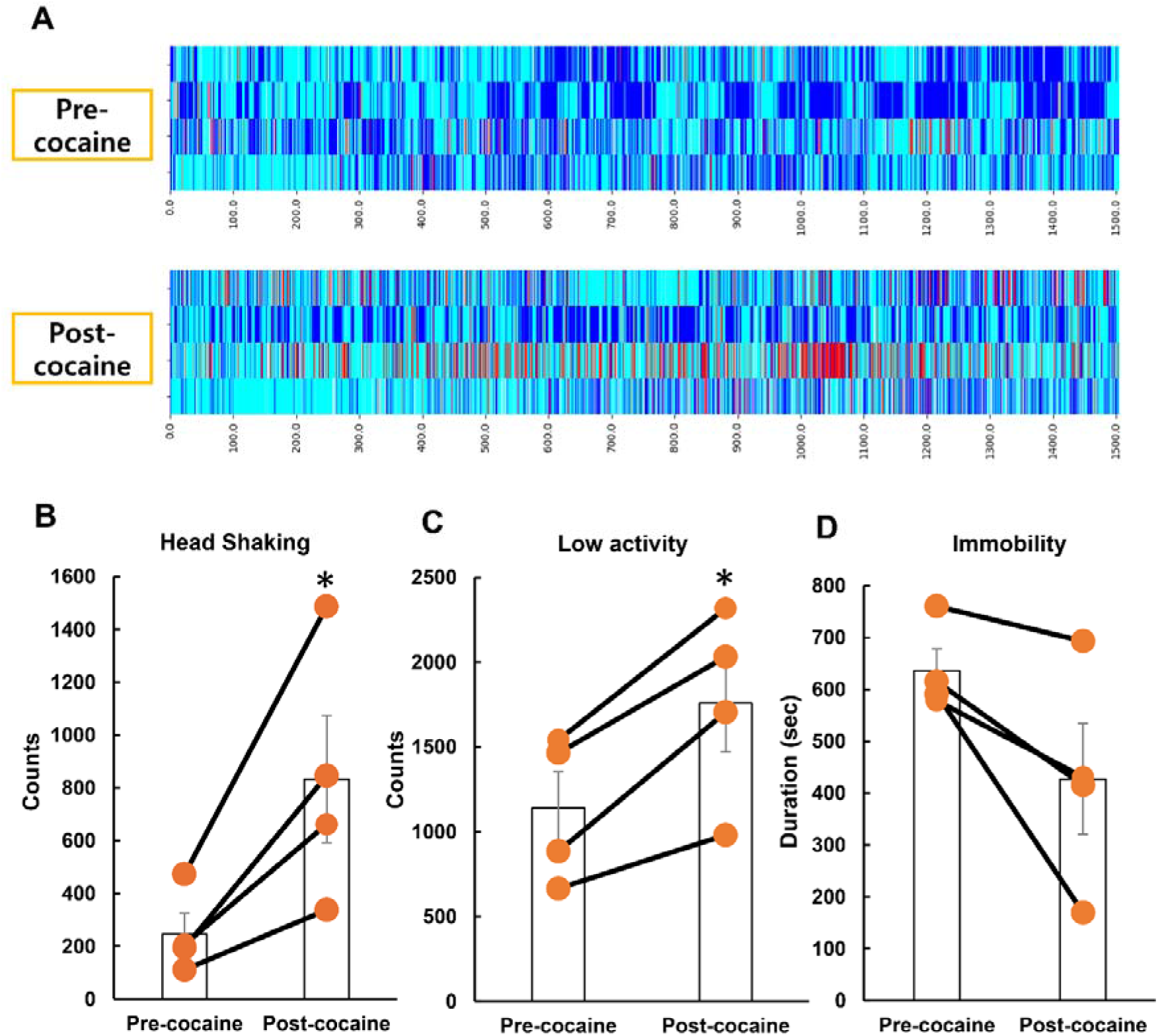
Artificial intelligence (AI)-based quantification of acute cocaine-induced behavioral changes in common marmosets. (A) LabGym-derived frame-wise raster plots show head shaking, low activity, and immobility during the 30-min pre- and post-cocaine periods; each row is one marmoset. Red indicates head shaking, cyan low activity, and blue immobility. (B–D) Head-shaking counts, low-activity counts, and immobility duration before and after acute cocaine (5 mg/kg, intraperitoneally). Orange circles show individual values, connecting lines pair observations, and bars show mean ± standard error of the mean (SEM) (*n* = 4). Statistical comparisons were performed using paired Student’s *t*-tests. Head-shaking counts and low-activity counts were significantly increased after cocaine administration, whereas the reduction in immobility duration was not statistically significant. \**P* < 0.05 versus pre-cocaine.

### Chronic cocaine administration alters functional brain connectivity

Across 15 predefined regions (Supplementary 1), resting-state functional connectivity decreased between the somatosensory cortex (SS) and posterior parietal cortex (PPC) at 1 month versus baseline (Fig. 2). At 10 months, connectivity decreased between the VLPFC and both the MPFC and motor and premotor cortex (MOT), but increased between the dorsolateral prefrontal cortex (DLPFC) and hippocampal formation (HipF). Repeated cocaine exposure therefore produced time-specific sensory, prefrontal, motor, and hippocampal connectivity changes.

**Fig. 2.**
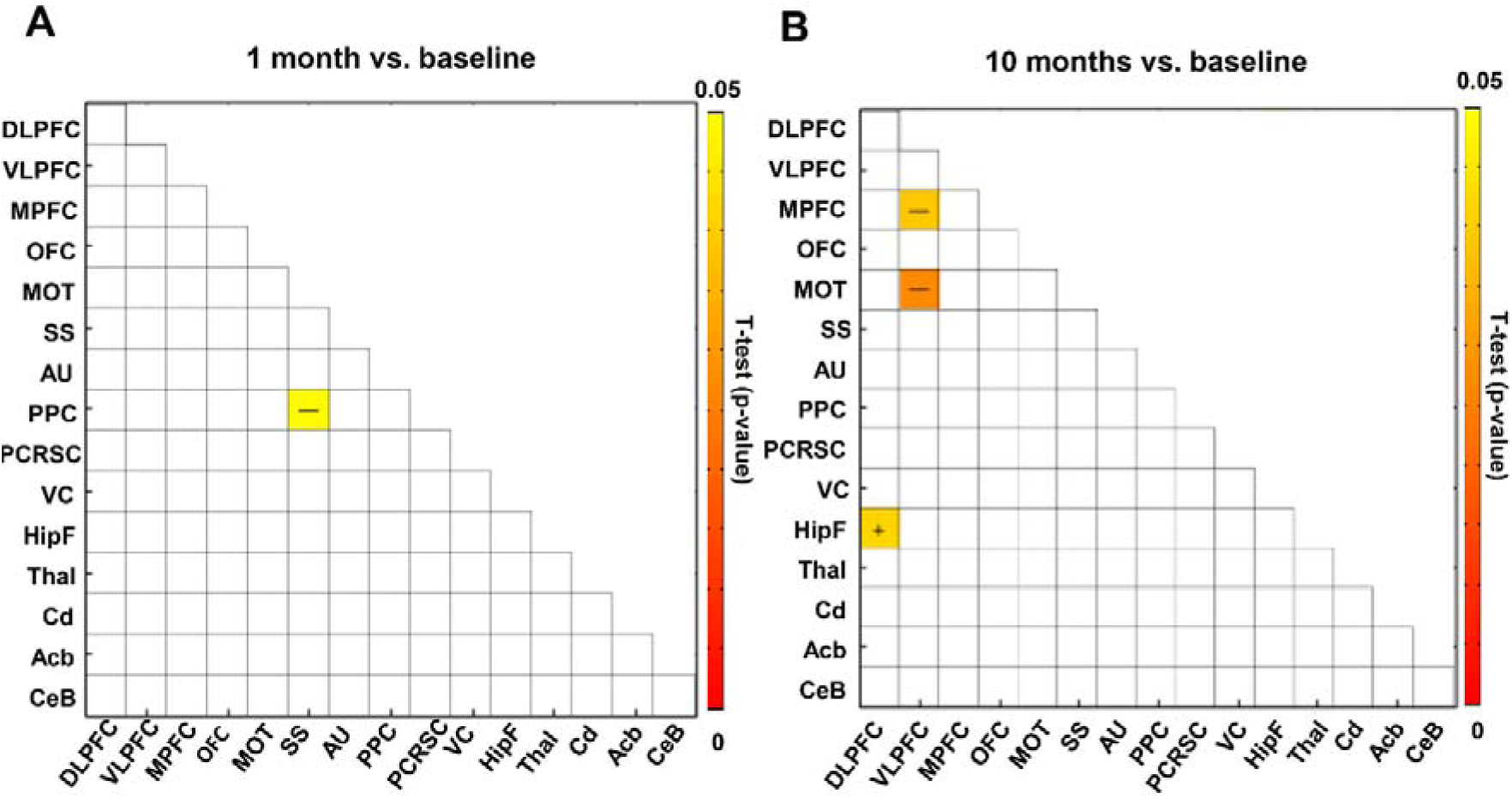
Resting-state functional connectivity after repeated cocaine administration. Region-of-interest (ROI)-based connectivity was evaluated across 15 cortical and subcortical regions in four common marmosets. (A) At 1 month versus baseline, somatosensory cortex (SS)-posterior parietal cortex (PPC) connectivity decreased. (B) At 10 months versus baseline, ventrolateral prefrontal cortex (VLPFC) connectivity with medial prefrontal cortex (MPFC) and motor and premotor cortex (MOT) decreased, whereas dorsolateral prefrontal cortex (DLPFC)-hippocampal formation (HipF) connectivity increased. Colored cells indicate paired Student’s *t*-test differences (*P* < 0.05, *n* = 4); lower *P* values are red. Plus and minus signs indicate increased and decreased connectivity, respectively.

### Repeated cocaine administration reduces white matter integrity in the splenium of the corpus callosum

Time significantly affected FA and AD in the SCC (FA, Greenhouse–Geisser-corrected *P* = 0.038; AD, *P* = 0.036; *n* = 4). Holm-adjusted comparisons showed lower SCC FA and AD at 10 months than baseline (FA, *P* = 0.012; AD, *P* = 0.011) (Fig. 3), whereas neither measure changed significantly in the genu (GCC). Repeated cocaine exposure thus produced region-specific SCC microstructural changes.

**Fig. 3.**
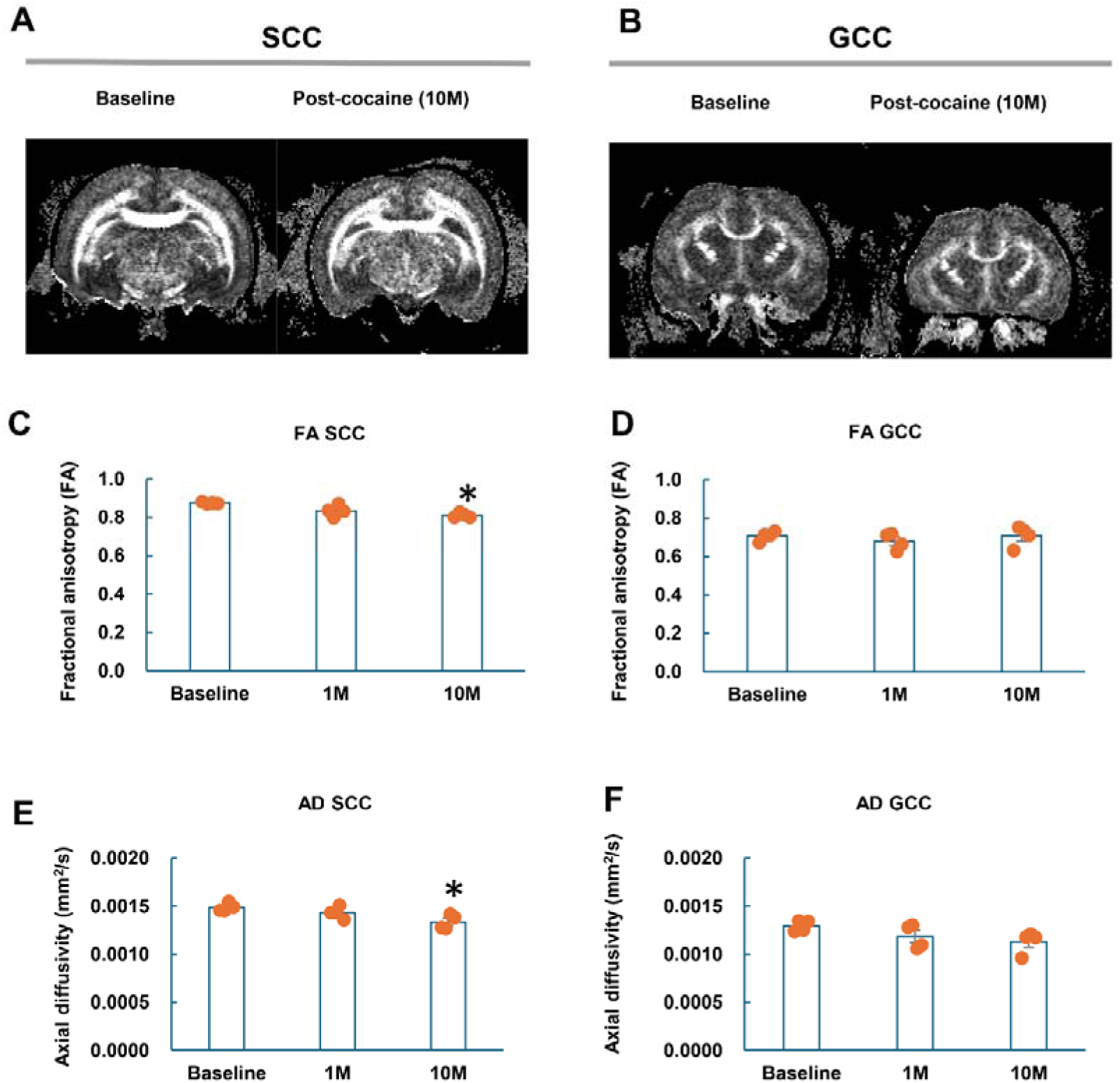
Repeated cocaine administration reduces FA and AD in the splenium of the corpus callosum. (A, B) Representative coronal DTI images of the SCC and GCC at baseline and 10 months after cocaine administration. (C–F) Quantification of fractional anisotropy (FA) and axial diffusivity (AD) in the SCC and GCC at baseline and at 1 and 10 months after cocaine administration. Significant effects of time were detected for FA and AD in the SCC. Holm-adjusted paired comparisons showed significant reductions at 10 months relative to baseline. No significant changes were observed in the GCC. Data are presented as the mean ± SEM with individual values (*n* = 4). \**P* < 0.05 versus baseline.

### Cross-species analysis identifies DPYSL2 as an axon-associated cocaine-responsive protein

Cross-species analysis compared our published human drug-user nucleus accumbens RNA-sequencing data (9) with proteomics of pooled marmoset cerebral cortex and underlying white matter. Gene Ontology cellular-component analysis identified 139 axon-associated human genes and 10 marmoset proteins; DPYSL2 and DNM3 overlapped, and DPYSL2 was selected for validation (Fig. 4A). In pooled proteomics, U3DL21 appeared only in control (normalized cocaine-to-control ratio, 0.0046), whereas U3F2W7 appeared only after cocaine (Supplementary Data 5), preventing inference of overall DPYSL2 direction. Western blotting of individual control (*n* = 3) and cocaine-treated (*n* = 3) samples showed significantly reduced DPYSL2 after chronic cocaine (Fig. 4B).

**Fig. 4.**
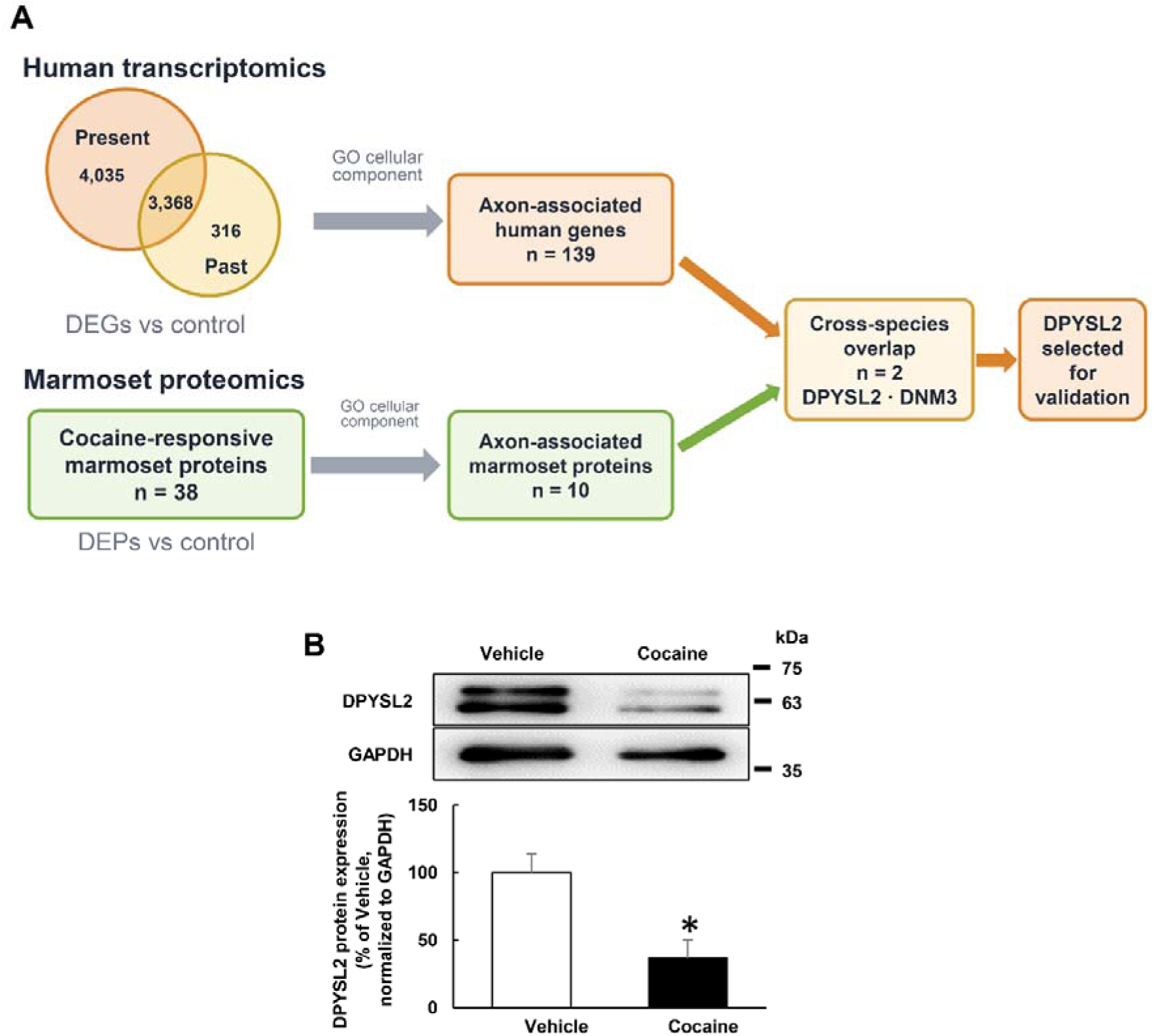
Cross-species identification and validation of dihydropyrimidinase-like 2 (DPYSL2) as an axon-associated cocaine-responsive protein. (A) Published human drug-user nucleus accumbens RNA-sequencing data (9) were compared with proteomics of pooled control and cocaine-treated marmoset cerebral cortex and underlying white matter. Gene Ontology cellular-component analysis identified 139 axon-associated human genes and 10 marmoset proteins; DPYSL2 and dynamin 3 (DNM3) overlapped, and DPYSL2 was selected for validation. (B) Representative Western blot and DPYSL2 quantification in control and 10-month cocaine-treated marmosets. Mean ± standard error of the mean (SEM) (*n* = 3 per group); unpaired Student’s *t*-test; \**P* < 0.05 versus control.

### CRYAB is expressed in corpus callosum oligodendrocytes and is increased after chronic cocaine exposure

CRYAB immunoreactivity colocalized with O4-positive oligodendrocytes in the marmoset corpus callosum (Fig. 5A). Western blotting of individual control (*n* = 3) and 10-month cocaine-treated (*n* = 3) samples showed increased CRYAB after chronic exposure (Fig. 5B), indicating a callosal oligodendrocyte protein response.

**Fig. 5.**
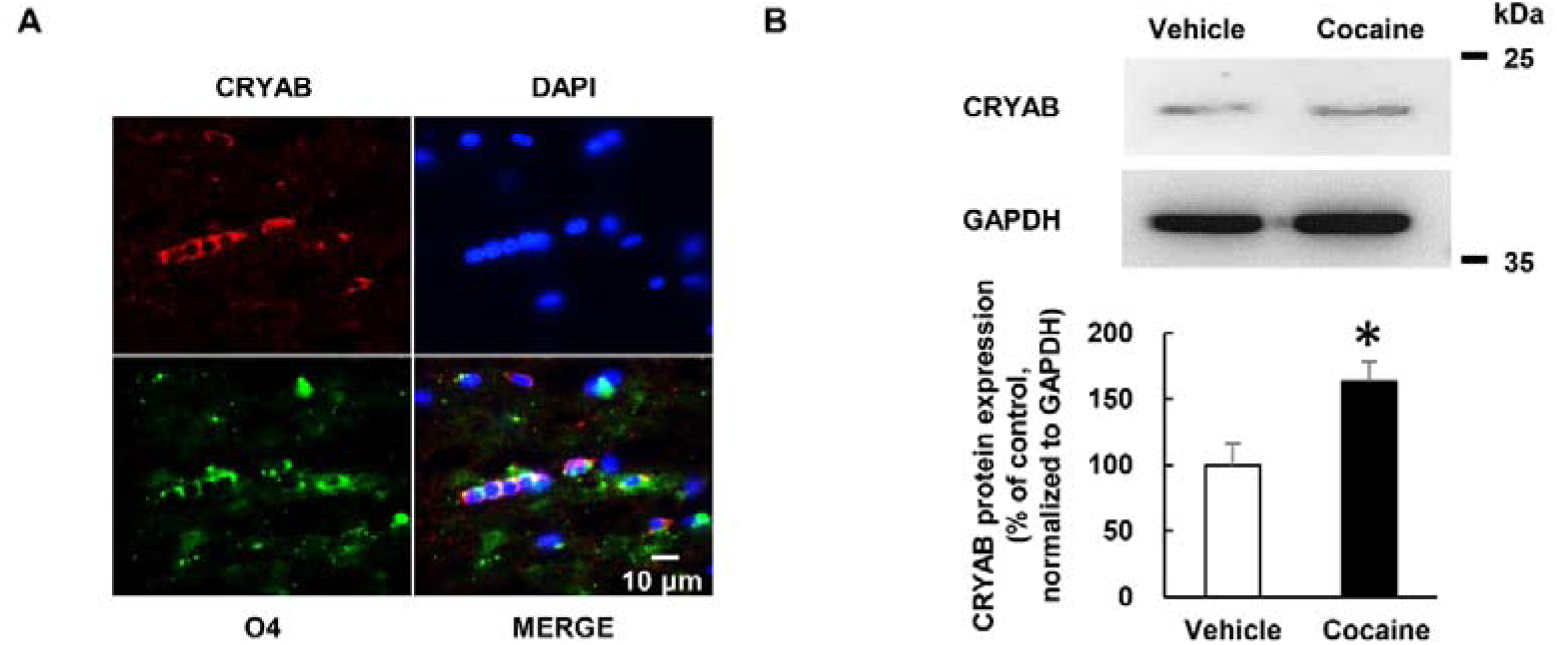
Crystallin alpha B (CRYAB) localization and expression in the marmoset corpus callosum after chronic cocaine exposure. (A) Representative immunofluorescence images show CRYAB (red), oligodendrocyte marker O4 (green), and 4′,6-diamidino-2-phenylindole (DAPI)-stained nuclei (blue). The merged image shows CRYAB in O4-positive oligodendrocytes. Scale bar, 10 μm. (B) Representative Western blot and CRYAB quantification in control and 10-month cocaine-treated marmosets. Mean ± standard error of the mean (SEM) (*n* = 3 per group); unpaired Student’s *t*-test; \**P* < 0.05 versus control.

### Loss of CRYAB is associated with reduced corpus callosum thickness and depression-related behavioral changes in mice

Brain morphology and behavior were assessed in *Cryab* knockout (KO) mice. At Bregma +1.1, +0.8, and +0.5 mm, Klüver–Barrera-stained sections showed thinner corpus callosa than in WT mice (Fig. 6A). Separate KO cohorts showed longer forced-swim immobility (WT, *n* = 6; KO, *n* = 6; Fig. 6B) and less locomotor activity (WT, *n* = 7; KO, *n* = 9; Fig. 6C). These findings associate CRYAB with callosal structure and depression-related behavior.

**Fig. 6.**
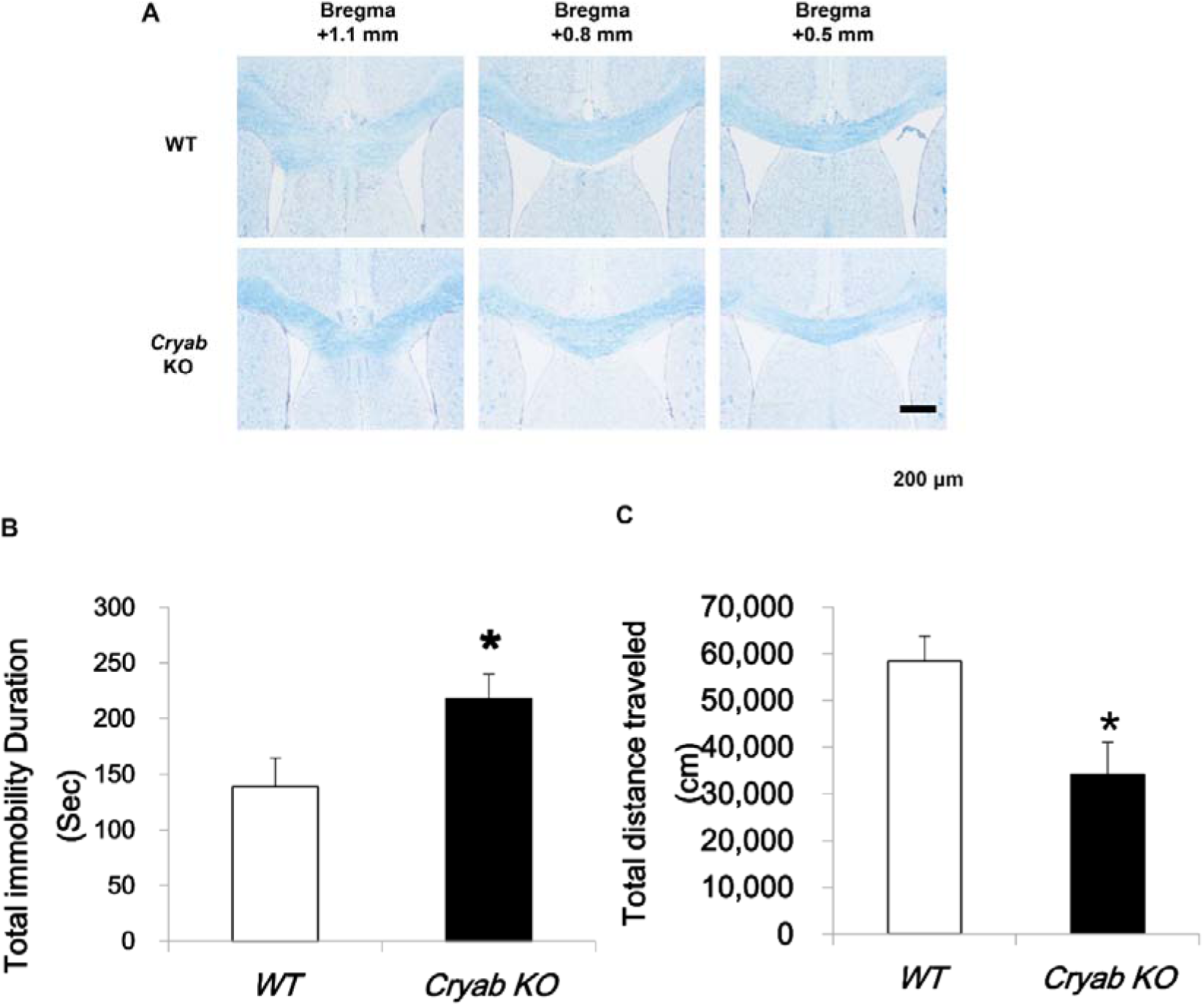
Effects of *Cryab* deletion on corpus callosum morphology and depression-related behavior in mice. (A) Representative Klüver–Barrera-stained coronal sections from wild-type (WT) and *Cryab* knockout (KO) mice at Bregma +1.1, +0.8, and +0.5 mm show reduced callosal thickness in KO mice. Scale bar, 200 μm. (B) Forced-swim immobility in WT and KO mice (*n* = 6 per group). (C) Locomotor distance in WT (*n* = 7) and KO (*n* = 9) mice. Separate cohorts were used in (B) and (C). Mean ± standard error of the mean (SEM); unpaired Student’s *t*-test; \**P* < 0.05 versus WT.

### Plasma cocaine concentrations

Cocaine was detected in all three cocaine-treated proteomics marmosets 4 h after the final dose. The mean plasma concentration was 26.3 ng/mL (86.3 nM; range, 14.8–45.8 ng/mL) (Table 1); Supplementary 3 provides representative chromatograms. These data confirmed systemic cocaine exposure.

**Table 1.** Concentration of cocaine in the plasma of marmosets.

| Measurement | Marmoset #1 | Marmoset #2 | Marmoset #3 | Mean |
| --- | --- | --- | --- | --- |
| Cocaine (ng/mL) | 45.8 | 18.4 | 14.8 | 26.3 |
| Cocaine (nM) | 150 | 60 | 49 | 86.3 |

Blood samples were collected 4 h after the final cocaine administration. Values represent individual measurements from three cocaine-treated marmosets.

## Discussion

Psychostimulants can induce stereotyped behaviors in animal models (1). Here, acute cocaine caused rapid, repetitive lateral head movements in all four marmosets, which LabGym distinguished from low activity and immobility. Repeated exposure then altered resting-state functional connectivity: somatosensory-posterior parietal connectivity decreased at 1 month; ventrolateral-medial prefrontal and ventrolateral-motor connectivity decreased at 10 months; and dorsolateral prefrontal-hippocampal connectivity increased at 10 months. Comparable functional changes occur in cocaine users (2,5) and nonhuman primates after long-term self-administration (3), which also produces structural and cognitive changes in rhesus macaques (4). Thus, acute behavior preceded connectivity changes and later structural abnormalities. Significant reductions in splenial FA and AD emerged at 10 months, while the genu was unchanged. Because the corpus callosum integrates bilateral cortical networks, early connectivity changes may reflect altered callosal communication. Reduced FA indicates altered white matter organization, and reduced AD can indicate axonal injury; similar abnormalities occur in chronic cocaine use (6). The splenium may therefore be more sensitive than the genu to prolonged exposure.

Cross-species analysis compared our published human drug-user nucleus accumbens RNA-sequencing data (9) with marmoset brain proteomics. Gene Ontology analysis found 139 axon-associated human genes and 10 marmoset proteins, with DPYSL2 and DNM3 shared. Opposing pooled DPYSL2 accessions—U3DL21 only in control (normalized cocaine-to-control ratio, 0.0046) and U3F2W7 only after cocaine—prevented directional inference from proteomics alone. Western blots of individual samples showed significantly reduced DPYSL2 after chronic cocaine.

DPYSL2, also called CRMP2, supports axon formation and microtubule assembly (16,17); its reduction may contribute to lower splenial FA and AD. Conversely, chronic cocaine increased CRYAB in O4-positive callosal oligodendrocytes. Our previous study also linked oligodendrocytic CRYAB to cocaine-related behavior (9). Because CRYAB protects cells during stress and demyelination (18), this increase may respond to white matter injury. *Cryab* knockout mice had thinner corpus callosa, greater forced-swim immobility, and less locomotor activity, supporting an association between CRYAB, callosal integrity, and depression-related behavioral changes.

Interpretation is limited by the small longitudinal cohort and absence of a time-matched untreated group. Pooled proteomics served only as an initial screen, followed by DPYSL2 and CRYAB validation in individual samples. Because resting-state functional connectivity does not directly measure callosal function, its relationship to later SCC changes requires further study.

Acute cocaine-induced head shaking preceded functional connectivity changes and later reductions in splenial FA and AD. Reduced DPYSL2 may relate to axonal injury, whereas increased CRYAB may respond to white matter stress; the mouse findings further associate CRYAB with corpus callosum integrity.

## Supporting information

Supplementary Data 5 - Marmoset brain proteomics

## Acknowledgments

This work was supported by the Ministry of Food and Drug Safety (20182MFDS425), the Bio&Medical Technology Development Program of the National Research Foundation (NRF) funded by the Korean government (MSIT) (No. RS-2024-00440787), and NRF Grant funded by MSIT (No. RS-2025-02273102).

