## Supplementary Information for "Acute head shaking precedes chronic corpus callosum deficits during repeated cocaine exposure in common marmosets"

**Supplementary 1. Regions of interest used for resting-state functional connectivity analysis**

Serial coronal sections of the customized common marmoset EPI template showing the 15 predefined regions of interest used for ROI-based functional connectivity analysis. The regions included the dorsolateral prefrontal cortex (DLPFC), ventrolateral prefrontal cortex (VLPFC), medial prefrontal cortex (MPFC), orbitofrontal cortex (OFC), motor and premotor cortex (MOT), somatosensory cortex (SS), auditory cortex (AU), posterior parietal cortex (PPC), posterior cingulate and retrosplenial cortex (PCRSC), visual cortex (VC), hippocampal formation (HipF), thalamus (Thal), caudate nucleus (Cd), accumbens nucleus (Acb), and cerebellum (CeB).

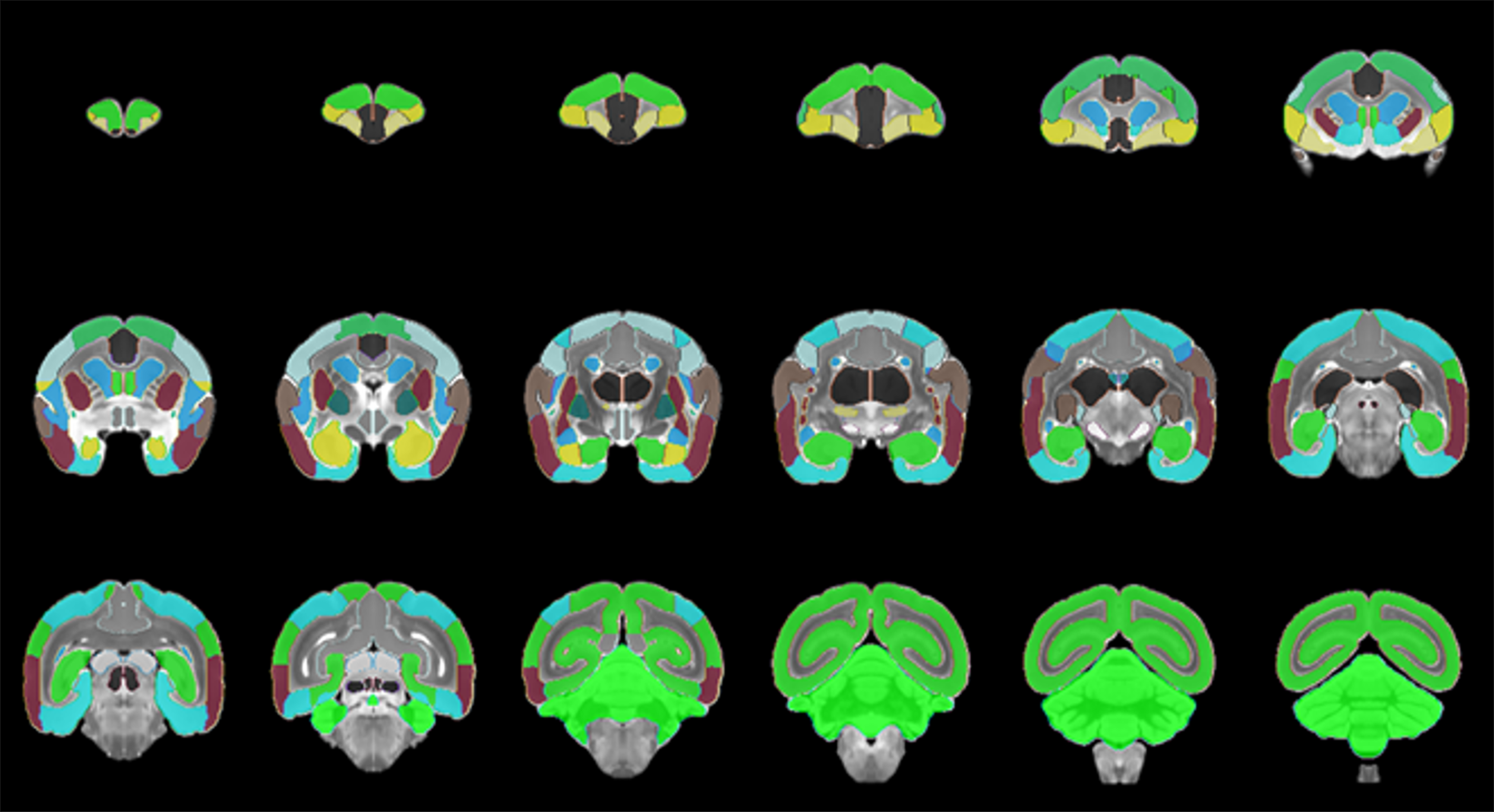

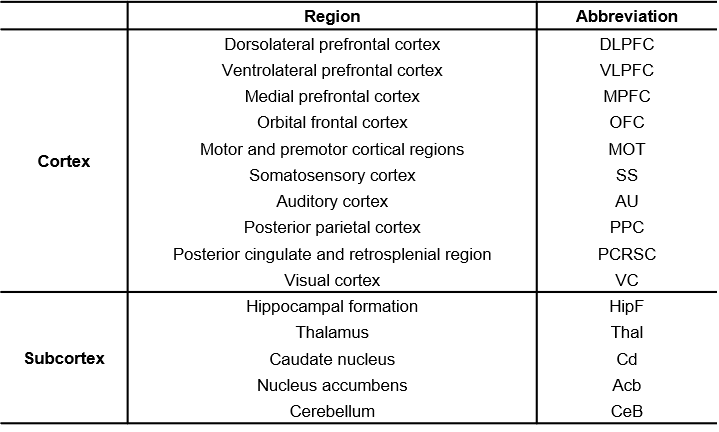

**Supplementary 2. LC–MS/MS acquisition parameters used for proteomic analysis.**

| **Nano LC gradient** | | | | **Full MS condition** | |
| --- | --- | --- | --- | --- | --- |
| **Time**  **(min)** | **Mobile**  **phase A (%)** | **Mobile**  **phase B (%)** | **Flow rate**  **(nL/min)** | Resolution | 70,000 |
| 0 | 95 | 5 | 300 | Scan range | 350–2,000 m/z |
| 10 | 95 | 5 |  | Maximum IT | 120 ms |
| 50 | 40 | 60 |  | Polarity | Positive |
| 53 | 5 | 95 |  | **MS/MS condition** | |
| 57 | 5 | 95 |  | Resolution | 17,500 |
| 60 | 95 | 5 |  | AGC | 5.00E+05 |
| 70 | 95 | 5 |  | Isolation width | 1.2 m/z |
| **Column temperature** | | | 50 ℃ | Top N | 20 |
|  |  |  |  | NCE (%) | 25 |
| **Sample temperature** | | | 4 ℃ | Maximum IT | 80 ms |
|  |  |  |  | Dynamic Exc. | 30 s |

LC-MS/MS, liquid chromatography–tandem mass spectrometry; AGC, automatic gain control; NCE, normalized collision energy

**Supplementary 3. LC-MS/MS Analysis for Cocaine**

Cocaine concentrations in marmoset plasma were quantified using an API 4000 triple-quadrupole mass spectrometer coupled to an Agilent 1100 HPLC system. Data acquisition and peak integration were performed using Analyst version 1.6.3 software. Chromatographic separation was performed using a Kinetex C18 100 Å column (2.1 × 50 mm, 2.6 μm).

The mobile phases consisted of 0.1% formic acid in distilled water as mobile phase A and methanol as mobile phase B. The flow rate was 0.2 mL/min, the column temperature was maintained at 40°C, and the injection volume was 10 μL. The gradient conditions are presented as follows:

| **Time (min)** | **A (%)** | **B (%)** |
| --- | --- | --- |
| **0.0** | 90 | 10 |
| **1.5** | 90 | 10 |
| **3.0** | 5 | 95 |
| **5.0** | 5 | 95 |
| **8.0** | 90 | 10 |
| **15.0** | 90 | 10 |

MS/MS analysis was performed using electrospray ionization in multiple reaction monitoring mode with positive polarity. The MRM transitions were selected based on previously reported methods and optimized for cocaine. The optimized MRM parameters are presented as follows:

| **Analyte** | **Q1** | **Q3** | **DP (V)** | **CE (V)** | **CXP (V)** |
| --- | --- | --- | --- | --- | --- |
| **Cocaine** | 304.5 | 182.1 | 56 | 27 | 12 |

DP, declustering potential; CE, collision energy; CXP, collision cell exit potential; MRM, multiple reaction monitoring.

Representative MRM chromatograms of plasma samples collected (A) vehicle control and (B) cocaine. Cocaine was monitored using the transition *m/z* 304.5 → 182.1. A cocaine peak was observed in the post-cocaine plasma sample at a retention time of approximately 6.18 min.

**A**

**Intensity (cps)**

**Time (min)**

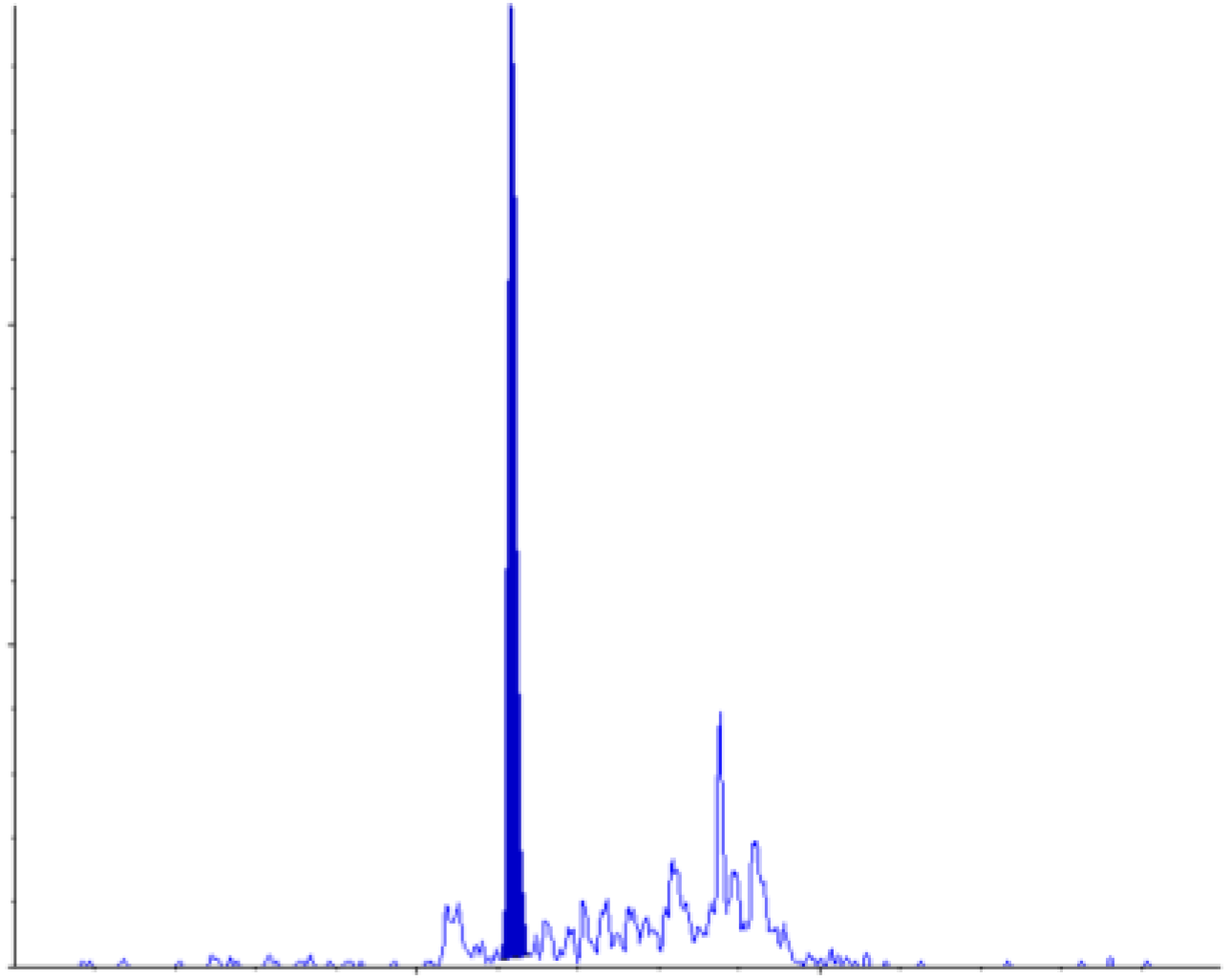

**6.18**

**1,000**

**500**

**0**

**5**

**10**

**8.76**

**5.51**

**Intensity (cps)**

**Time (min)**

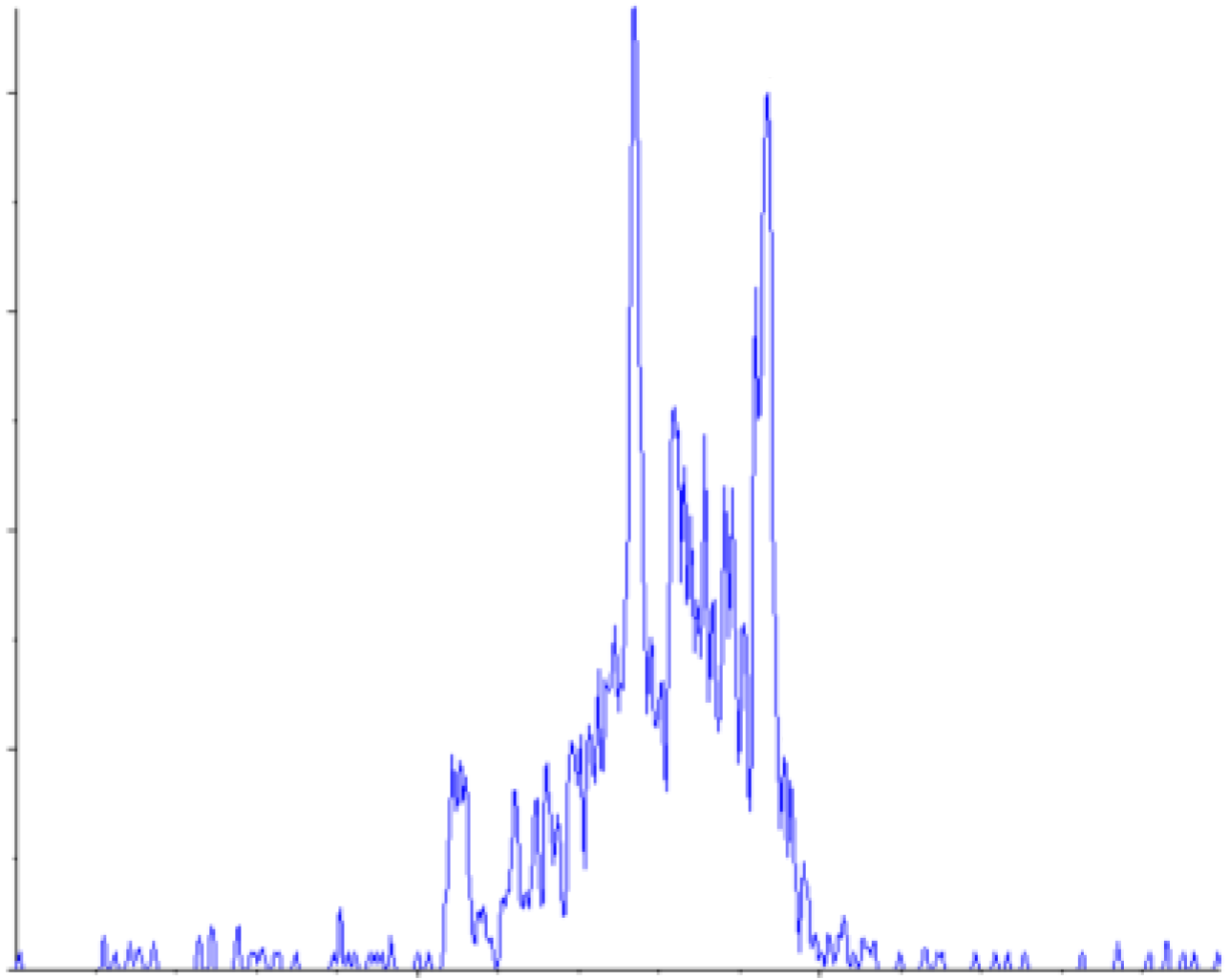

**300**

**400**

**200**

**0**

**7.69**

**4.04**

**7.45**

**9.55**

**9.35**

**5**

**10**

**100**

**B**
