## Supplementary Data 4 - Video for "Acute head shaking precedes chronic corpus callosum deficits during repeated cocaine exposure in common marmosets"

### Slide 1
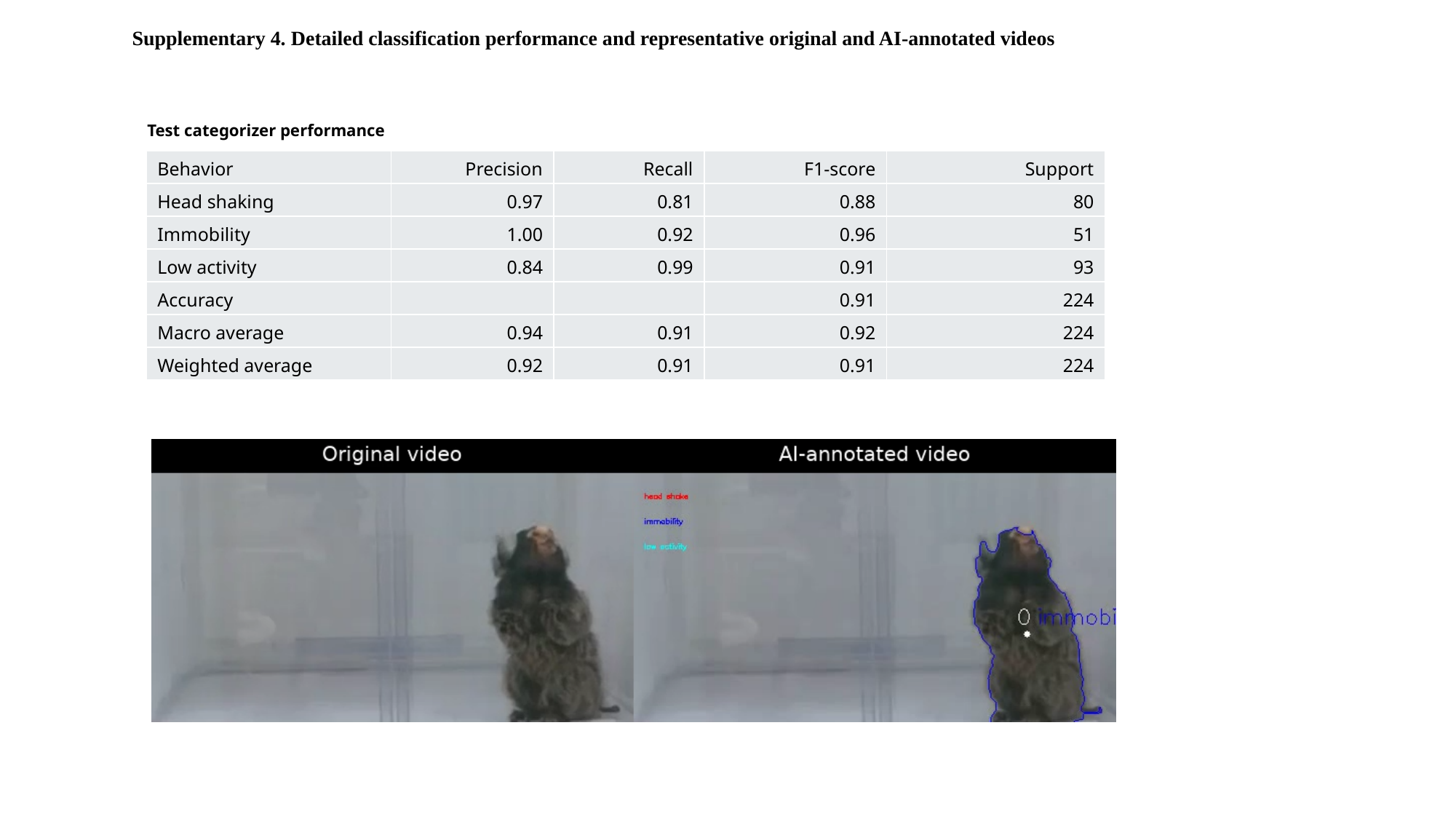

Supplementary 4. Detailed classification performance and representative original and AI-annotated videos
Test categorizer performance
| Behavior | Precision | Recall | F1-score | Support |
| --- | --- | --- | --- | --- |
| Head shaking | 0.97 | 0.81 | 0.88 | 80 |
| Immobility | 1.00 | 0.92 | 0.96 | 51 |
| Low activity | 0.84 | 0.99 | 0.91 | 93 |
| Accuracy | | | 0.91 | 224 |
| Macro average | 0.94 | 0.91 | 0.92 | 224 |
| Weighted average | 0.92 | 0.91 | 0.91 | 224 |
